# Control of odor sensation by light and cryptochrome in the *Drosophila* antenna

**DOI:** 10.1101/2024.12.12.628259

**Authors:** Dhananjay Thakur, Sydney Hunt, Tiffany Tsou, Miles Petty, Jason M. Rodriguez, Craig Montell

## Abstract

Olfaction is a sense employed by insects to differentiate safe from harmful food options, evaluate potential mates, and identify oviposition sites. Here, we found that the fruit fly, *Drosophila melanogaster*, responds differently to a set of repulsive odors depending on ambient light conditions. Ultraviolet (UV) or blue light reduces the behavioral aversion and electrophysiological responses of olfactory receptor neurons (ORNs) to certain repellent odors, such as benzaldehyde. We found that *cryptochrome* (*cry*) is strongly expressed in the antennal support cells that lie adjacent to ORNs, and mutation of *cry* eliminates the light-dependent reduction in aversion. Thus, these data demonstrate that support cells in an olfactory organ serve a sensory function as light receptor cells. It has been shown that light activation of Cry creates reactive oxygen species (ROS), and ROS activates the TRPA1 channel. We demonstrate that the TRPA1-C isoform is expressed and required in ORNs for benzaldehyde repulsion, and that TRPA1-C is activated *in vitro* by benzaldehyde. Overexpression of dual oxygenase, which generates hydrogen peroxide, reduced the aversion under dark conditions. Our data support the model that light-dependent creation of hydrogen peroxide persistently activates TRPA1-C. Consequently, the channel is no longer effectively activated by benzaldehyde. Since flies sleep much more at night, and begin feeding at dawn, we propose that the light-induced reduction in aversion to certain odors provides a mechanism to lower the barrier to feeding following the transition from night to day.

## Introduction

Chemosensation includes the senses of taste and smell, and it is known that taste is greatly influenced by multiple types of sensory stimuli. In humans, the perception of sweetness and bitterness is affected by food temperature, food texture, sound and external light.^1–8^ The receptor mechanisms underlying how these sensory features impact taste have been characterized using model organisms, ranging from flies^9–13^ to mice.^14^ In the case of olfaction, some people associate certain odors with specific colors.^15,16^

Due to the association of colors with certain odors, in principle, environmental light could impact odor acuity or valence. However, this has not been documented. To address this question, we elected to focus on the fruit fly, *Drosophila melanogaster*, since the olfactory system, as well as the behavioral and electrophysiological responses to a wide range of odorants have been studied extensively in flies.^17^ Moreover, the possibility that light could directly affect olfaction is particularly plausible in insects such as the fruit fly, since the two olfactory organs, the antennae and the maxillary palps, are external.^18^ While the 3^rd^ segment of the antenna contains a high density of olfactory receptor neurons (ORNs), other parts of the antennal structure, namely the Johnston’s organ and arista, are sensitive to stimuli such as humidity^19–21^, sound^22,23^ wind^24^, gravity^25^, and changes in temperature.^26^ The possible integration of odor detection with sensation of other physical parameters is under active investigation. Smell acuity, for example, is known to be sensitive to ambient temperature,^27,28^ and changes during the circadian rhythm.^29^

*Drosophila* antennae are situated on the dorsal surface of the fly and are directly and constantly exposed to light. Although they are known to express photoreceptive proteins such as the ultraviolet (UV) and blue light sensitive rhodopsins^30^, and cryptochrome (Cry),^31^ it is not known if the antennae sense and respond directly to ambient light, or if light has an impact on the odor response.

Here, we discovered that ambient, short-wavelength light reduces the aversion to a select set of noxious odorants, such as benzaldehyde. The light is detected directly by olfactory sensilla in the antenna, and this sensation depends on Cry, which is known to be expressed in central pacemaker neurons in the brain, where it contributes to light-entrainment of circadian rhythms.^32–35^ Surprisingly, Cry is required in support cells rather than ORNs in the sensilla. Our work supports the model that light activation of Cry results in induction of reactive oxygen species, which causes persistent activation and subsequent refractoriness of the TRPA1 channel in ORNs to responsiveness to benzaldehyde. This work highlights a role for light in modifying the olfactory response, which does so through light reception in a new class of light receptor cells present in an olfactory organ.

## Results

### DART2 assay for olfactory avoidance of repellents

To assay whether olfactory avoidance in *Drosophila* is impacted by light, we modified a two-way choice assay (direct airborne repellent test, DART)^36^ consisting of a tube capped at one end with a repellent and the solvent that was used to solubilize and dilute the repellent at the other end, with lights directed from above (Figure 1A). To ensure that there was no contact between the flies and the chemicals during the assay, the caps included meshed chambers, creating a separation of 8 mm between the flies and the chemicals. We transferred flies to the tube, allowed them to acclimate for 20 minutes without any odorants, added the odorants, and then recorded their positions over the course of 60 minutes. To optimize the assay, we first focused on characterizing olfactory behavior in the dark under near-infrared illumination with a video camera (Figure 1A).

**Figure 1.**
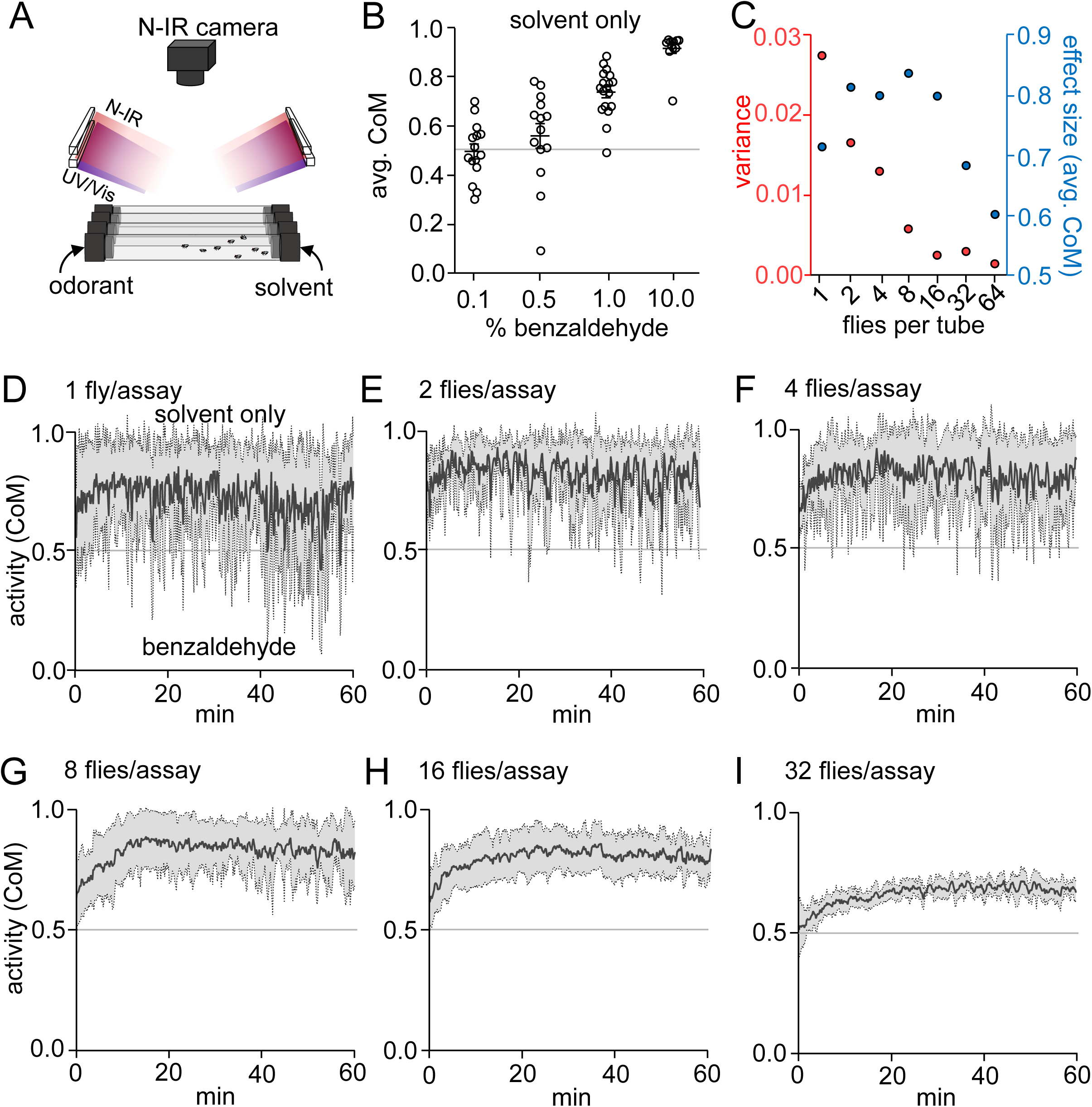
DART2 assay for assaying olfactory behavior. (A) A two-way choice DART2 assay set up using glass tubes with meshed chambers carrying odorants or solvents. Flies were imaged using reflected near-infrared (N-IR) light. Images were captured at the rate of 1 frame/sec and the center of mass (CoM) of the flies was analyzed at 1 frame/10 sec. (B) Average CoM of control flies in response to varying concentrations of BA from 20– 60 min of the duration of the assay under N-IR illumination only. (C) Calibration of the number of flies used per tube. Summary of variance and effect size plotted against number of flies per tube, which were exposed to 1% BA. Each population test was repeated 5 – 6 times. (D) 1 fly per tube. (E) 2 flies per tube. (F) 4 flies per tube. (G) 8 flies per tube. (H) 16 flies per tube. (I) 32 flies per tube. See also Figure S1.

We tracked the responses of the flies by calculating the transverse component of their center of mass (CoM) per frame and normalizing it to the length of the tube. We found that the CoM provided the most precise quantitative alignment with the visually observed distribution of flies in the assay tube. The most extreme avoidance or attraction to a chemical, as seen when the flies cluster at one end or the other (Figure S1A), produces a CoM of 1.0 or 0 (Figure S1B). If the flies distributed homogeneously throughout the tube (Figure S1A), this would indicate no aversion or attraction to the chemical and result in a CoM of 0.5 (Figure S1B). Other sample distributions indicating less than the most severe repulsion (e.g. right bias, right cluster or slight right bias; Figure S1A) produce CoM >0.5 – <1.0 (Figure S1B), while less than the most severe attraction (e.g. left bias, left cluster or slight left bias; Figure S1A), result in CoM >0 – <0.5 (Figure S1B). In contrast to the CoM metric, measuring the position of the flies in terms of avoidance index (Figure S1C) could not discriminate between subtle differences in the distribution of the flies within the tubes. Therefore, we used CoM as the measure of odor responses throughout this study.

Benzaldehyde (BA) is a naturally occurring compound synthesized in plants and the odor is repellent to *Drosophila*.^37^ The flies were indifferent to 0.1% or 0.5% BA, while increasing the BA to just 1% elicited strong avoidance (Figure 1B). Elevating the BA to 10% caused extreme avoidance, with the flies clustering near the right side of the DART2 tube (Figure 1B).

In order to calibrate the variability of the response as a function of the number of flies in the DART2 assay, we tested the behavioral response to 1% BA with different numbers of flies ranging from 1 to 64. We found that there was considerable variation in the response over time if the assays included only 1, 2, or 4 flies (Figures 1C–F). In close concordance with previous work that shows that collective behavior produces more robust responses to aversive cues,^38^ we found that increasing the number of flies in the assay (≥8) resulted in less variability in the response to BA (Figures 1C, G–I and S1D). In assays with 8 and 16 flies, the CoM moved increasingly away from BA over the first 20 minutes, reaching a steady-state of aversion that was maintained for the remainder of the 60-minute-long assay (Figures 1G and H). Including greater numbers of flies in the assay (32 or 64) caused slight decreases in the average CoM relative to the assays with 16 (Figures 1I and S1D) as some of the flies were forced into the side of the tube closer to the repellent due to overcrowding. In order to obtain minimal variance and maximum effect size (Figure 1C), we elected to use 8-16 flies per tube for the remaining analyses. Next we tested whether there was sexual dimorphism in the response to BA. We found that the responses were indistinguishable when we tested males only, females only, or a mix of both sexes (Figure S1E). Therefore, we used a mix of males and females in each tube for all further analyses.

### UV and blue illumination reduce aversion to benzaldehyde

To determine whether exposure to light impacted the responses to BA, we exposed flies to white light illumination and performed DART2 assays. We found that in the presence of white light, the avoidance of 1% BA was significantly reduced (Figure 2A). We then tested the impact of different wavelengths of light and found that ultraviolet light (UV; 365 nm) or blue light (450-480 nm), but not green light (520 nm) or red light (625 nm), suppressed the repulsion to BA (Figure 2B). The aversion was decreased significantly when the flies were exposed to ≥20 μW/cm^2^ of UV or ≥18 μW/cm^2^ blue illumination (Figures 2C and D). We assayed the impact of 100 μW/cm^2^ UV light on repulsion by different concentrations of BA ranging from 0.1–10%. While the response to 1.0% benzaldehyde was suppressed by UV, there were no effects of UV on lower BA levels (0.1% and 0.5%), and the highest concentration of BA (10%) was so aversive (CoM = 0.93 ±0.01) that the UV light had no impact (Figure 2E).

**Figure 2.**
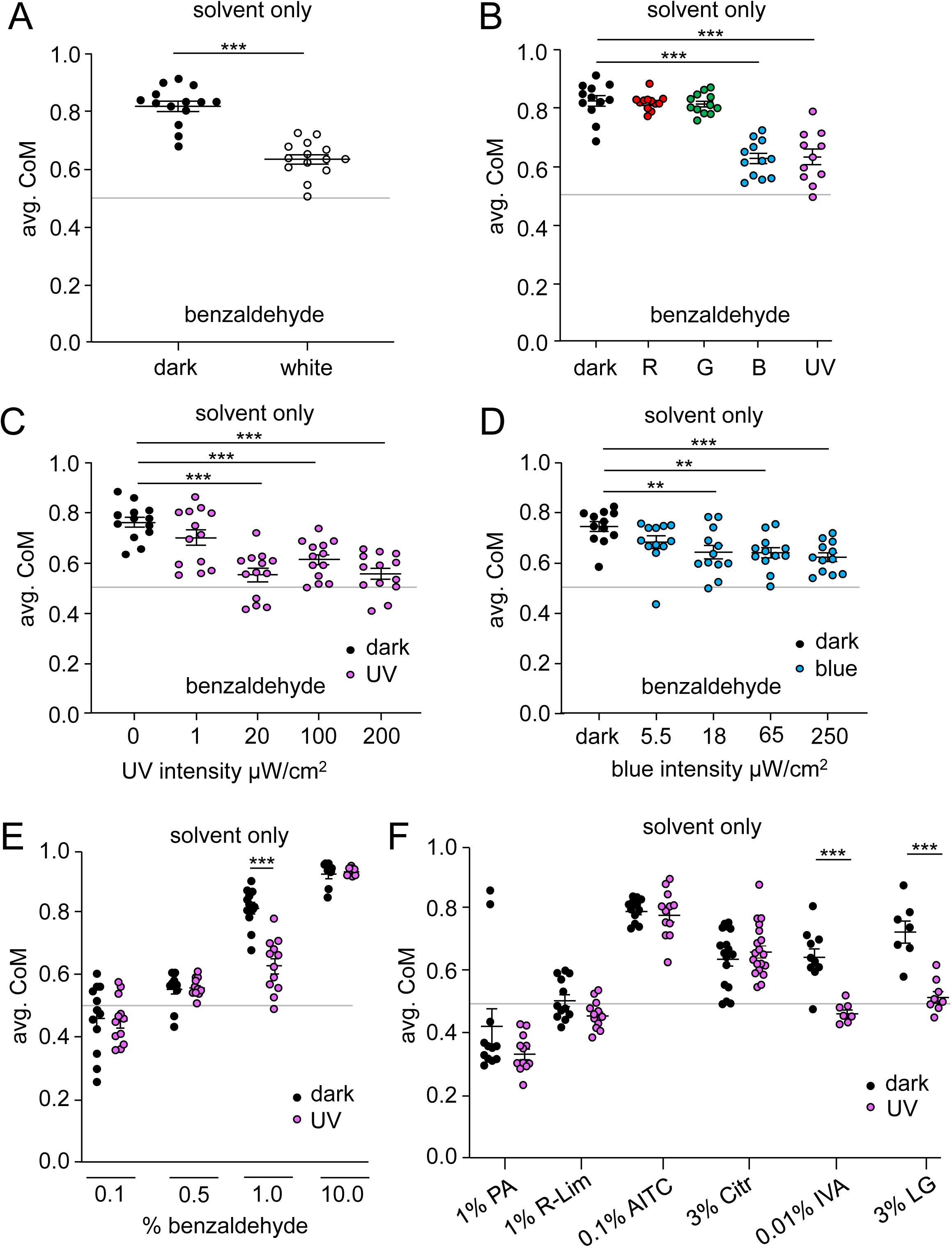
Impact of light on olfactory aversion. (A) Average CoM of control flies in response to 1% BA under dark conditions (0–0.01 μW/cm^2^) or under white light (250 μW/cm^2^). White light was supplied by simultaneously applying red, green, blue and UV LEDs. n = 14. Mann-Whitney U test. (B) Average CoM of control flies in response to 1% BA with red, green, blue or UV illumination, with the light intensity at 100 μW/cm^2^ for each wavelength. n = 12. ANOVA followed by Dunn’s multiple comparisons test. (C) Average CoM of control flies in response to 1% BA as a function of UV intensity. n = 13. ANOVA followed by Dunn’s multiple comparisons test. (D) Average CoM of control flies in response to 1% BA as a function of intensity of blue light (450 nm). n = 12. ANOVA followed by Dunn’s multiple comparisons test. (E) Average CoM of control flies in response to varying concentrations of BA under UV illumination at 100 μW/cm^2^. n = 12. Mann-Whitney U test. (F) Average CoM of control flies in response to 1% propionic acid (PA), 1% R-limonene (R-lim), 0.1% allyl isothiocyanate (AITC), 3% citronellal (Citr), 0.01% isovaleric acid (IVA) and 3% lemongrass (LG) under UV illumination at 100 μW/cm^2^. n = 12. Mann-Whitney U test. Error bars indicate mean ± SEMs. ** p < 0.01, *** p < 0.001.

To determine if UV light modified the responses to other odorants, we tested AITC (allyl isothiocyanate), which is a potent noxious odorant to flies,^39^ propionic acid (a constituent of fly food), limonene (a neutral odor), citronellal, which is a previously described repellent,^36^ isovaleric acid (a product of fermentation) and lemongrass, a widely used insect repellent.^40^ The responses of the flies were not significantly different under UV and dark conditions to propionic acid, limonene, AITC, and citronellal (Figure 2F). However, similar to benzaldehyde, the aversion was significantly reduced when flies were tested with isovaleric acid and lemongrass (Figure 2F).

### ORN responses are suppressed by light, via a rhodopsin independent mechanism

To test if light directly modulates the response of the ORNs, we performed electroantennograms (EAGs), which are field recordings performed by placing a recording electrode on the antenna. We exposed flies to 1% BA either in the dark or during 5 minutes of blue light. Upon exposure to blue light, the response to BA was attenuated to 41.6 ±6.2% of the response in the dark (Figures 3A and B), showing that light directly affects ORN activity in the antenna. These results suggest that there is a light sensor in the antenna that serves to modulate the olfactory response.

**Figure 3.**
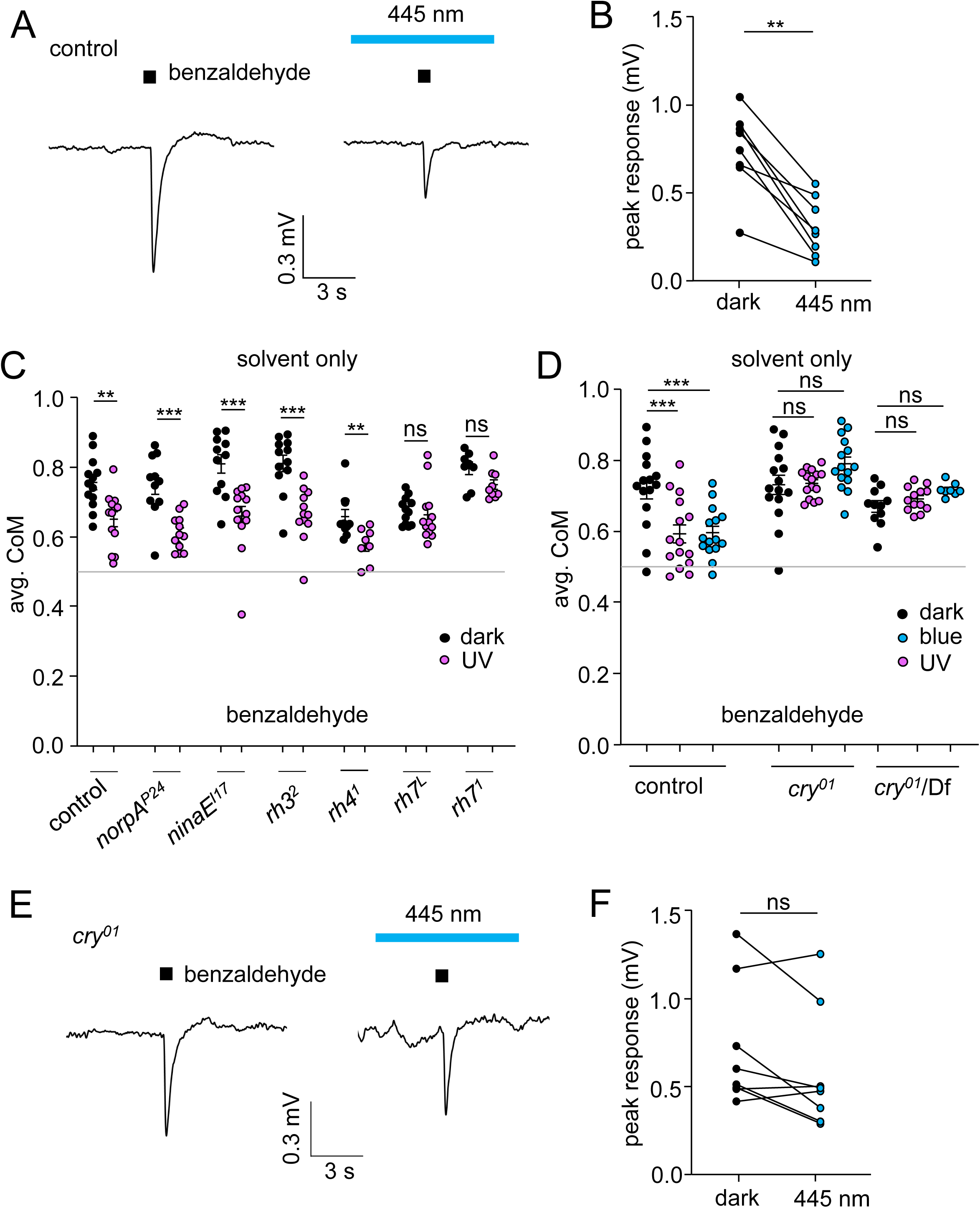
Direct effect of light on antennal odor responses. (A) EAG responses in control flies exposed to 1% BA in the dark (left) or under blue light for 5 min (right). (B) Peak EAG responses. n= 8. Paired Wilcoxon test. (C) Average position of CoM of control, *norpA^P24^, rh1* (*ninaE*)*, rh3, rh4* and *rh7* null mutants from 20 - 60 min during the assay under UV illumination at 100 μW/cm^2^. n = 11 – 13. Mann-Whitney U test. (D) Average CoM of control, *cry*^01^ and *cry*^01^/Df in response to 1% BA from 20 - 60 min during the assay under UV illumination (100 μW/cm^2^) or blue illumination (250 μW/cm^2^). n = 8 – 13. ANOVA followed by Dunn’s multiple comparisons test. (E) EAG responses in control and *cry*^01^ flies exposed to 1% BA in the dark (left) or under blue light for 5 min (right). (F) Summary of peak EAG responses. n= 8. Data were analyzed using paired Wilcoxon test. ns = not significant Error bars indicate mean ± SEMs. ** p < 0.01. *** p < 0.001. See also Figure S2.

To identify the light receptor required for suppressing the repulsion to BA in the antenna, we tested the requirements for rhodopsins that are activated by UV and blue light. Flies encode seven rhodopsins, four of which are effectively activated by UV and/or blue light (Rh1, Rh3, Rh4 and Rh7).^41–44^ Six of the rhodopsins (Rh1-Rh6) engage a trimeric G-protein that couples to a phospholipase Cβ (PLC) encoded by the *norpA* gene,^18^ while Rh7 signaling is mediated through PLC21C.^42^ We found that flies lacking *norpA* showed similar aversion to BA under UV light as control flies (Figure 3C), indicating that rhodopsins that engage this G-protein-activated phospholipase Cβ are not the receptors enabling the light-dependent suppression of BA repulsion. Consistent with this finding, mutations affecting *rh1* (*ninaE*), *rh3* or *rh4* did not alter the impact of light on the aversion to BA (Figure 3C). In contrast, flies lacking *rh7* showed no difference in aversion between dark and UV conditions. However, we did not detect *rh7*-promoter-driven GFP staining in the antenna (Figure S2), suggesting that there exists another light sensor in the antenna that modifies ORN responses to BA.

### Cryptochrome required for UV and blue light suppression of olfactory avoidance

Cryptochrome (Cry), which detects UV/blue light in *Drosophila* central pacemaker neurons in the brain,^45,46^ is also expressed in the antenna.^31^ Therefore, Cry is an excellent candidate sensor for functioning in the light-dependent modification of the response to BA in the antenna. We discovered that a null mutation in *cry* (*cry*^01^) eliminated the suppression of the olfactory avoidance to BA by UV or blue light (Figure 3D). We observed a similar phenotype when we placed the *cry*^01^ mutation *in trans* with a deficiency that uncovered the *cry* locus (Figure 3D). Thus, we conclude that Cry is required for the UV- and blue-induced suppression of BA repulsion.

To determine whether *cry* impacts the electrical response to BA in the antenna, we performed EAGs. As mentioned above, light suppressed the BA response in wild-type antennas (Figures 3A and B). In contrast, in *cry*^01^ flies the ability of blue light to reduce the peak EAG amplitude was diminished <20% (81.5 ±8.4% of the peak in the dark), demonstrating that *cry* provides a large fraction of the inhibition in odor response by light in the antenna (Figures 3E and F).

### *cry* is expressed and required in non-neuronal cells in olfactory sensilla

The two main olfactory organs are the bilaterally symmetrical 3^rd^ antennal segments and the maxillary palps.^18^ These organs are decorated with bristles, referred to as olfactory sensilla, which harbor one to four ORNs, and three types of accessory cells. These include the trichogen (shaft), tormogen (socket), and thecogen (sheath).^47^ To visualize the cells that express *cry*, we took advantage of a *cry* reporter (*cry-GAL4*)^48^ to drive expression of *UAS-mCD8::GFP*. The reporter stained cells in both the 3^rd^ antennal segment and the maxillary palp (Figures 4 and S3).

**Figure 4.**
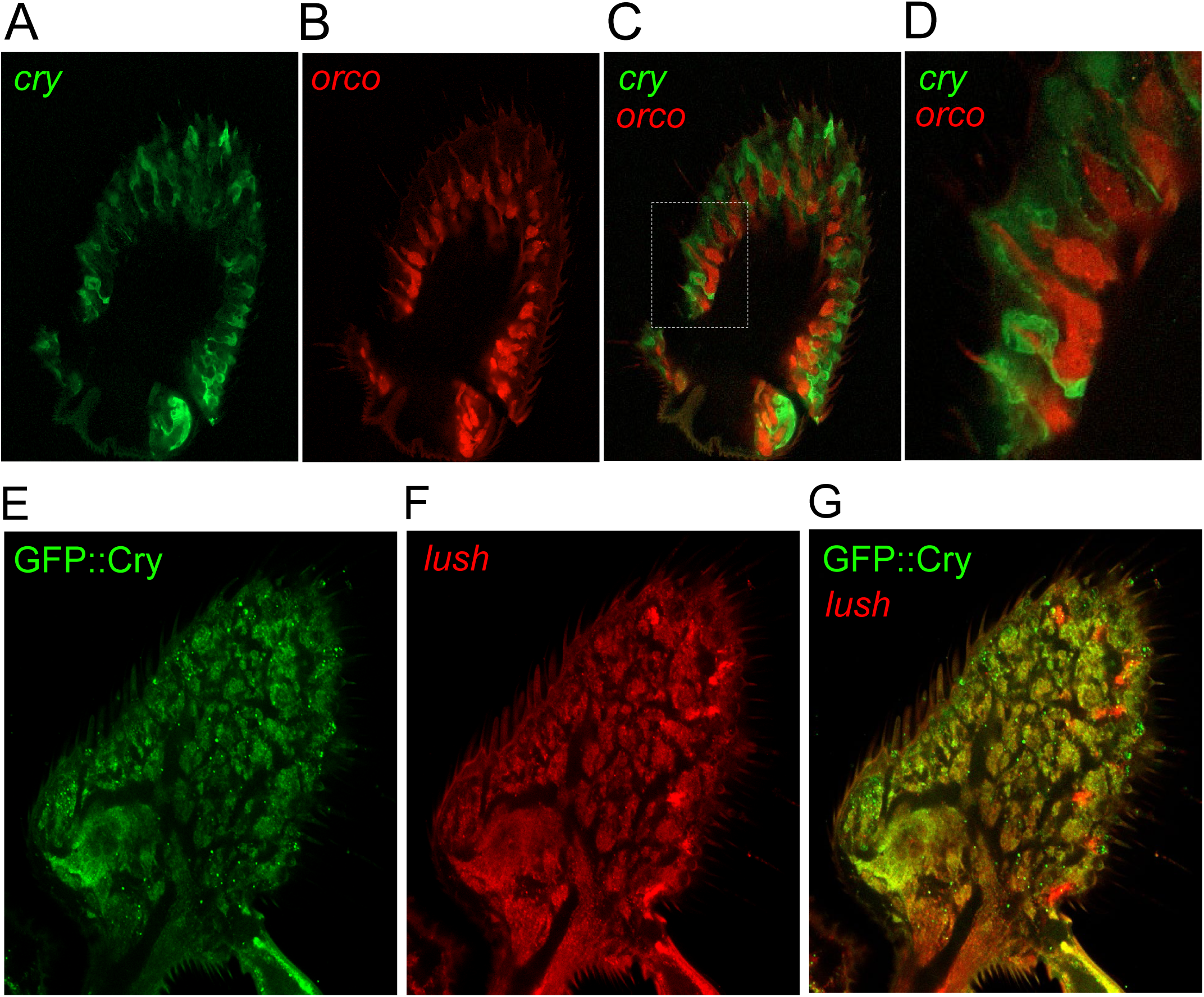
Immunostained of olfactory organs. (A) Antennal cross-section showing *UAS-mCD8::GFP* expression driven by *cry-GAL4* and stained with an anti-GFP antibody. (B) Same sample as (A) showing expression of *RFP* driven directly by the *orco* promoter (*orco-RFP*) and stained with anti-RFP. (C) Merge of (A) and (B). (D) Magnified section, marked in (C) indicated by white dotted rectangle. (E) Antennal cross-section showing expression of GFP-tagged Cry (GFP::Cry) stained with anti-GFP. (F) Same sample as (E) showing expression of *UAS-DsRed* driven by the *lush-GAL4* and stained with anti-DsRed. (G) Overlay of (E) and (F). See also Figures S3, S4 and S5.

Cry is expressed in central pacemaker neurons ^32,34,49^, suggesting that it olfactory organs is it also expressed in neurons. To address whether *cry* is expressed in ORNs, we performed double-labeling experiments using the *cry*-*GAL4* in combination with a marker specific for ORNs, *orco-RFP,* which expresses *RFP* under control of the *orco* promoter.^50^ Surprisingly, we did not detect co-staining with the *orco-RFP* either in the antenna (Figures 4A–D) or in the maxillary palp (Figures S3A–D). Moreover, there were no *cry*-positive axonal processes extending into any glomeruli in the antennal lobes in the brain (Figures S3E–G). Together, these data indicate that the *cry* reporter is not expressed in ORNs. To address whether Cry is expressed in support cells, we took advantage of the GFP::Cry fusion protein,^51^ which stained the same set of cells as the *cry-GAL4* (Figure S3H–J). We found that nearly all of the GFP::Cry positive cells in the antenna and maxillary palps were also stained with the *lush-GAL4* (Figures 4E–G and S3K–M), which labels a large population of support cells throughout the antenna and primarily seems to overlap with cells labeled by *ase5-GAL4*.^52^ These experimental observations align well with our analysis of transcript data from the FlyCellAtlas that is consistent with *cry* expression in support cells in olfactory sensilla (Figure S4).

To identify the types of support cells that express *cry*, we used *GAL4*s to drive expression in the tormogen (*ase5*), thecogen (*nompA*), and glia (*repo*). Consistent with the extensive overlap of the *cry* and *lush* reporters, we found that the GFP::Cry showed considerable overlap with all three cell types (Figure S5). To determine whether *cry* is sufficient in olfactory support cells, we performed rescue experiments using the *UAS-cry* and the *lush-GAL4*, and found that this recapitulated the reduced aversion to BA in the presence of UV light (Figure 5A).

**Figure 5.**
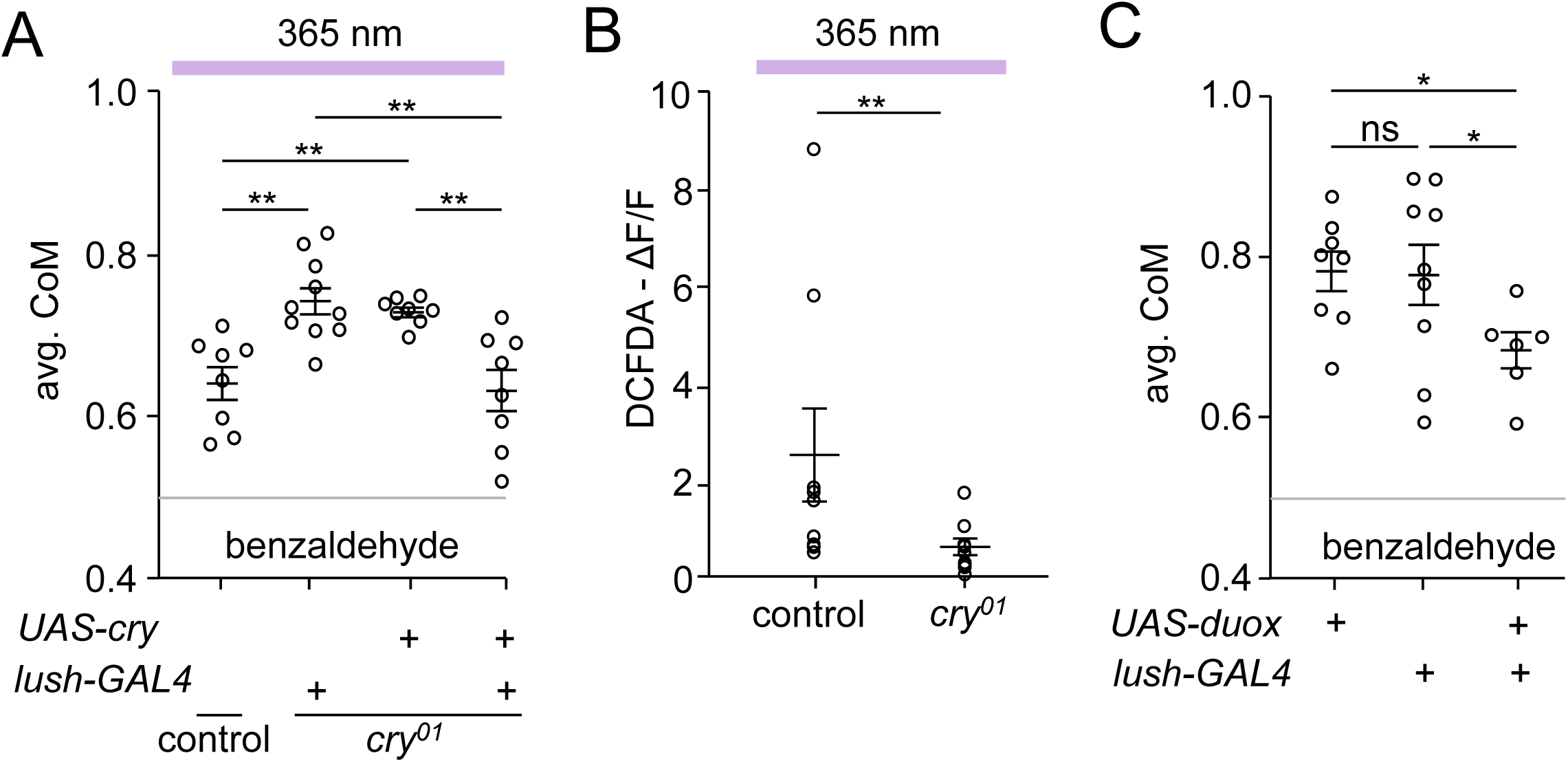
Characterizing the role of *cry* in the antenna. (A) Rescue of cry expression (*UAS-cry*) in lush expressing cells (*lush-GAL4*) shows similar reduction as controls in aversion to BA under UV illumination. n = 8-10. ANOVA followed by Dunn’s multiple comparisons test. (B) Levels of ROS generation in isolated antennae under UV stimulation, shown in terms of change in fluorescence intensity (ΔF/F) using DCFDA. n = 14 – 15. Student’s unpaired *t*-tests. (C) Induction of oxidative stress in support cells is sufficient to reduce aversion to BA. *H. sapiens UAS-duOx* was expressed using the *lush-GAL4*. n = 4-5. ANOVA followed by Dunn’s multiple comparisons test Error bars indicate mean ± SEMs. * p < 0.05. ** p < 0.01. *** p < 0.001. ns = not significant.

### Role of reactive oxygen species in modulating odor response

The surprising finding that Cry expression in support cells is sufficient for UV-induced suppression of BA repulsion raises the question as to the underlying mechanism. One possibility is that Cry in support cells leads to release of a diffusible product that impacts ORN function. Activation of Cry by short-wavelength light changes the oxidative state of the flavin FADH that is bound to Cry, resulting in the production of reactive oxygen species (ROS).^53^ The ROS could in principle affect the activity of the ORNs.

To test whether activation of Cry by UV light increases production of ROS in the antennae, we used the ROS sensor, 2′,7′-dichlorofluorescein diacetate (DCFDA), as described previously.^53^ We found that when we surgically removed antennae from control flies and incubated them with DCFDA, they showed a significant increase in fluorescence response after 1 second of UV exposure (Figure 5B). In contrast, the response was reduced in *cry*^01^ mutant antennae (Figure 5B). The residual fluorescence response was likely an effect of UV on other endogenous photosensitizers in the tissue.^54,55^ To determine whether induction of oxidative stress in *lush*- and *cry*- expressing olfactory support cells is sufficient to reduce aversion to BA, we used the *lush-GAL4* to express dual oxidase (*UAS-duox*), which encodes a protein that generates hydrogen peroxide. We found that this was sufficient to attenuate the repulsion to BA in the dark (Figure 5C).

### TRPA1-C is required for avoidance to benzaldehyde

The observation that *cry* is required for UV-induced ROS production in the antenna raises the question as to the molecular target for the ROS. ROS, including H_2_O_2_, which are produced by UV light, activate *Drosophila* TRPA1,^56,57^ and we have demonstrated previously that a *trpA1* reporter is expressed in ORNs in the 3^rd^ antennal segment.^36^ The *trpA1* gene is expressed as four primary isoforms (A-D).^36,58–60^ Both *trpA1-A* and *trpA1-B* use one promoter and *trpA1-C* and *trpA1-D* share an alternative promoter. To address which isoform-pair is expressed in ORNs, we used *GAL4* lines that we previously created that are specific to the *AB* isoforms (*trpA1-AB^GAL4^*) and the *CD* isoforms (*trpA1-CD^GAL4^*).^61^ The axons of ORNs extend into different glomeruli in the olfactory lobes. Thus, using *UAS-mCD8::GFP*, which labels the cell surfaces of cell bodies and neuronal processes, we could determine which driver is expressed in ORNs by examining labeling of the 58 glomeruli in each olfactory lobe.^62–65^ Using the *trpA1-AB*-*GAL4*, we did not detect significant staining of olfactory glomeruli (Figure 6A). In contrast, the *trpA1-CD-GAL4* labeled ≥13 glomeruli in each lobe (Figures 6B and S6). To determine whether the *trpA1-C* or the *trpA1-D* isoform is predominantly expressed in ORNs we employed the *trpA1-C-T2A-GAL4* and *trpA1-D-T2A-GAL4*s.^60^ We detected significant staining in the antennal lobe driven by the *trpA1-C-T2A-GAL4* but not by the *trpA1-D-T2A-GAL4* (Figures 6C and D).

**Figure 6.**
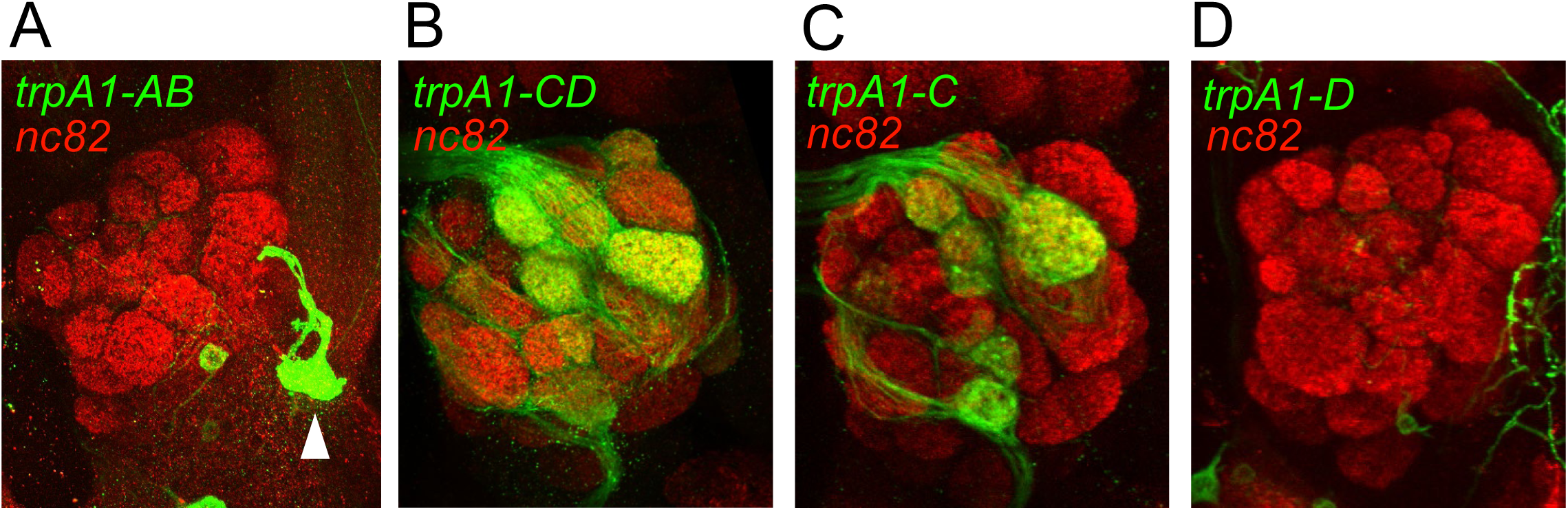
Examining staining in the antennal lobes showing neuronal projections expressing reporters for different *trpA1* isoforms. (A) Antennal lobe expressing *UAS-GFP* under the control of the *trpA1-AB-GAL4*, and stained with anti-GFP and anti-nc82. The white arrowhead indicates an anterior cell (AC) neuron.^77^ (B) Antennal lobe expressing *UAS-GFP* under the control of the *trpA1-CD-GAL4*, and stained with anti-GFP and anti-nc82. (C) Antennal lobe expressing *UAS-GFP* under the control of the *trpA1-C-T2A-GAL4*, and stained with anti-GFP and anti-nc82. (D) Antennal lobe expressing *UAS-GFP* under the control of the *trpA1-D-T2A-GAL4*, and stained with anti-GFP and anti-nc82. See also Figure S6.

To determine whether the *trpA1-C* isoform is sufficient for BA avoidance, we used two lines of flies. First, we analyzed *trpA1-C* knockin flies (*trpA1-C-KI*),^60^ which express *trpA1-C* in the absence of any other isoform, and found that the *trpA1-C-KI* flies exhibited BA avoidance similar to control flies (Figure 7A). In addition, we performed rescue experiments by expressing either the *UAS-trpA1-C* or the *UAS-trpA1-D* transgene in a *trpA1^1^* null mutant background under control of the *trpA1-CD-GAL4*. Expression of *trpA1-C* rescued the *trpA1* mutant phenotype, while there was only a small effect of expressing the *trpA1-D* isoform, which was not significant (Figure 7B). Furthermore, when we inactivated *trpA1-C* ORNs by expressing an inwardly rectifying K^+^ channel (*UAS-kir2.1*) under control of a *trpA1-C-T2A-GAL4*, the repulsion to 1% BA was reduced significantly (Figure 7C).

**Figure 7.**
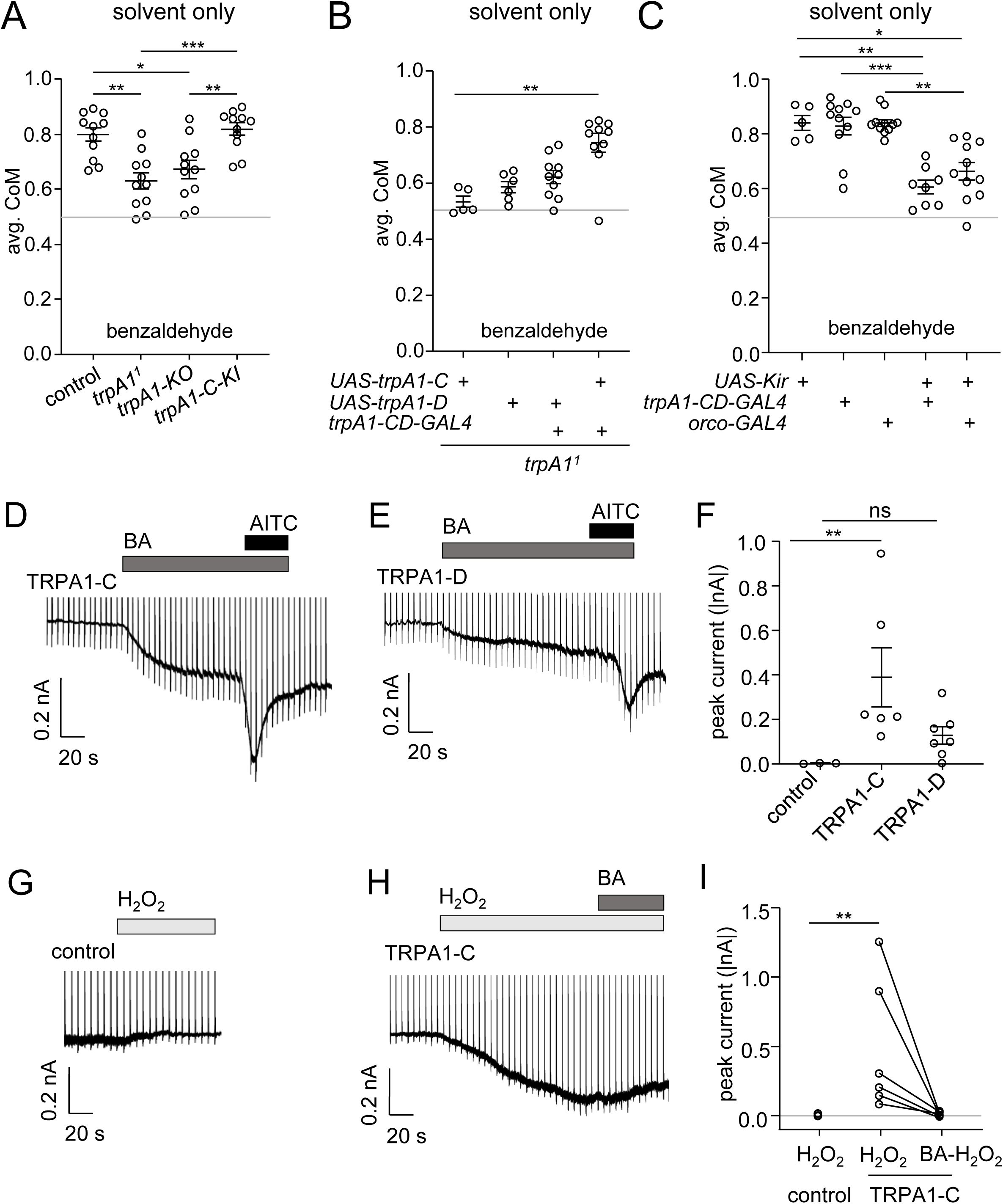
The role of TRPA1 in response to benzaldehyde and ROS. (A) Average CoM of control, *trpA1^1^, trpA-KO* and *trpA1-C-KI* flies in response to 1% BA from 20 - 60 min during the assay. n = 11. Data were analyzed by ANOVA followed by Dunn’s multiple comparisons test. (B) Average CoM of *trpA1^1^* flies expressing *UAS-TRPA1-C* or *UAS-TRPA1-D* driven by the *trpA1-CD-GAL4* in response to 1% BA from 20 - 60 min during the assay. (C) Average CoM of flies expressing *UAS-Kir2.1* driven by *trpA1-CD-GAL4* or the *orco- GAL4* in response to 1% BA from 20 - 60 min during the assay. n = 5 – 11. (D) Representative whole-cell currents from S2 cells expressing TRPA1-C in response to 0.1% BA and 0.1% AITC. (E) Representative whole-cell currents from S2 cells expressing TRPA1-D in response to 0.1% BA and 0.1% AITC. Low concentrations of BA and AITC were sufficient to activate large whole cell currents because TRPA1-C was overexpressed. (F) Summary of peak current responses to 0.1% BA from cells expressing only GFP (control), TRPA1C or TRPA1D. n = 3 – 7. Data were analyzed by ANOVA followed by Dunn’s multiple comparisons test. (G) Whole-cell representative currents from S2 cells expressing only GFP (control) in response to 100 µM H_2_O_2_. (H) Whole-cell representative currents from S2 cells expressing TRPA1-C in response to 100 µM H_2_O_2_ and 0.1% BA. (I) Summary of peak current responses to 0.1% BA from cells expressing either GFP only (control) or TRPA1-C. n = 4 – 6. Paired Wilcoxon test. Error bars indicate mean ± SEMs. * p < 0.05. ** p < 0.01. *** p < 0.001. ns = not significant.

To determine whether TRPA1-C is directly activated by BA, we expressed TRPA1-C in an insect cell line (S2 cells) and performed whole-cell recordings. We applied BA dissolved in the extracellular bath solution, and terminated each recording with an activator of TRPA1 channels, allyl isothiocyanate (AITC).^66–68^ We found that BA activated TRPA1-C with an average current of 389.1 ±132.6 pA (Figures 7D and F). The TRPA1-D isoform was also activated by BA, but the peak current was much smaller (128.1 ±39.1 pA; Figures 7E and F). Therefore, we conclude that the TRPA1-C channel is critical for conferring repulsion to BA.

Next, we asked whether exposing cells expressing TRPA1-C to H_2_O_2_ would prevent TRPA1-C from being activated by BA. We found that 100 μM H_2_O_2_ robustly activated TRPA1-C-dependent currents (Figures 7G-I). When we applied BA to the same cells that had been exposed to H_2_O_2_ for 2 minutes, we found that BA did not result in any further activation of TRPA1-C (Figures 7H-I).

## Discussion

In this work, we provide the first demonstration that light directly impacts odor detection. Specifically, we show that the flies’ avoidance of certain botanically-derived chemicals, such as BA, IVA, and lemongrass is reduced in the presence of light, relative to the dark. Flies sleep to a greater extent at night than during the day, especially compared to dawn and dusk. We suggest that the reduced avoidance in a light environment results because the flies become more active, and need to begin to forage after the onset of light.^69–72^ Therefore, the barrier to feeding is much higher in the dark, and should decrease in the light. Food options are complex mixtures of many chemicals, some of which are attractive on their own, while others are aversive. Therefore, in the light, when the barrier to feeding declines, we suggest that light decreases the flies’ aversion to some repulsive volatile chemicals in a potential food, thereby allowing them to land on the substrate and use contact chemosensation and other senses to further evaluate the food.

Surprisingly, the mechanism for the light-induced reduction in olfactory avoidance depends on light sensation in the antenna by Cry, which was originally identified due to its role in light detection in circadian pacemaker neurons in the fly’s brain.^32–35^ Also, unexpected is the observation that Cry functions in support cells in olfactory sensilla, rather than in neurons. The finding that Cry is the light sensor in the antenna provides an explanation for the salience of UV and blue light, rather than longer wavelengths, for suppressing olfactory aversion, since Cry is activated by these shorter wavelengths.^45,46^ Moreover, this work identifies support cells in olfactory sensilla as a new class of light receptor cell.

Four rhodopsins (Rh1, Rh3, Rh4 and Rh7) sense short wavelength light.^41–44^ However, mutations disrupting the *rh1* (*ninaE*), *rh3* and *rh4* genes do not diminish the suppression of olfactory repulsion by light. The Rh1, Rh3 and Rh4 rhodopsins couple to a PLCβ encoded by the *norpA* locus, and consistent with the findings with these mutants, a null mutation in *norpA* also does not reduce the light-induced suppression of the BA repulsion. However, disruption of *rh7* does diminish the effect of light. Nevertheless, we do not detect *rh7* reporter expression in the antenna, indicating that it does not contribute to the effect of light on the olfactory response in the antenna. Rh7 is expressed in many neurons in the brain.^42^ Therefore, its contribution to the light-induced suppression of olfactory repulsion may reflect a function in higher order neurons in the brain.

A key question concerns the mechanism through which expression of Cry in antennal support cells leads to suppression of olfactory avoidance in ORNs. It has been shown previously that UV/blue light activation of Cry leads to ROS production,^53^ and using a ROS sensor we found that short-wavelength light generated ROS in the antenna, which was reduced in the *cry* mutant background. Conversely, expression of DuOx in support cells, which promotes production of hydrogen peroxide, reduces BA repulsion in the absence of UV/blue light. Of note, *Drosophila* TRPA1 is activated by ROS,^56,57^ and we found that TRPA1-C is expressed in ORNs. Moreover, we showed that TRPA1-C contributes to olfactory repulsion of BA, and is directly activated by BA. We propose the model that ROS produced by light activation of Cry in support cells leads to desensitization of TRPA1-C in ORNs, which in turn attenuates the aversion to BA. These findings also highlight a role that support cells play in actively modulating the response to an odor. Although we were not able to maintain our whole-cell recordings sufficiently long to determine if H_2_O_2_ results in complete return to the baseline (desensitization), we found that within a couple of minutes of H_2_O_2_ exposure, the current began to decline, and was not enhanced upon application of BA.

According to our model, a simple prediction is that light would suppress the repulsion to all aversive compounds that activate TRPA1-C. However, light did not reduce the repulsion to AITC, even though it is a well-established activator of TRPA1 in *Drosophila*,^36,39^ and of homologs in other animals.^66–68^ This raises the possibility that TRPA1-C is not the only receptor for AITC. Consistent with this possibility, repulsion of volatile AITC depends on an odorant receptor, OR42a.^73^ Moreover, since volatile AITC is highly toxic to *Drosophila*,^73^ a reduction in repulsion to this chemical would reduce survival, and so the observation that repulsion to AITC is not reduced by light is consistent with a likely requirement to strongly avoid AITC at all times. Citronellal is a relatively weak activator of *Drosophila* TRPA1, yet light does not attenuate the repulsion to this chemical either. This may be because citronellal is also detected through olfactory receptors (ORs).^36^ Benzaldehyde can also be sensed in part through ORs.^74^ We suggest that may account for our findings that the olfactory repulsion to BA is reduced but not eliminated in the *trpA1* mutant, and that UV/blue light attenuates rather than completely suppresses the olfactory repulsion to BA.

In conclusion, we discovered that light can modify olfactory reception and behavior, and this occurs through direct light activation of cells in olfactory sensilla. Moreover, the light is detected by support cells, demonstrating that these cells are a class of non-neuronal light receptors, which then affect the olfactory response in ORNs.

## Materials and Methods

### Fly stocks and husbandry

Flies were reared at 25 °C on normal cornmeal food under 12 hr light/12 hr dark cycles. The control flies carried *w^+^* and the 2^nd^ and 3^rd^ chromosomes were from *w^1118^* (BDSC #5905). All mutants were outcrossed to *w^1118^* for five generations. The following lines were used (BDSC stock numbers): *trpA1^1^* (26504), *trpA1-AB^GAL4^* (67131), *trpA1-CD^GAL4^* (67133)*, cry-GAL4* (24514), *cry^Df^* (27922)*, cry*^01^ {Dolezelova et al. 2007}, *orco-RFP* (63045), *orco-GAL4* (23292), *ase5-GAL4* (93029), *nompA-GAL4* (605353), *repo-GAL4* (7415), *UAS-hDuOx* (78412), *UAS-Cry* (86263), *UAS-mCD8::GFP* (5137), *10xUAS-IVS-mCD8::RFP*, *13xLexAop2-mCD8::GFP* (32229), *norpA^P24^*(9048), *ninaE^I17^*(5701), *rh3^2^* {Li et al, 2020}, *rh4^1^* {Vasiliauskas et al, 2011}, *rh7^1^* {Ni et al, 2017}, *rh7^L^ (rh7-LexA)* {Meyerhof et al, 2024}, *UAS-dsRed*.^75^ Y. Xiang provided *trpA1-KO, trpA1-C-KI, trpA1-C-T2A-GAL4* and *trpA1-D-T2A-GAL4*.^60^ W.D. Tracey provided *UAS-trpA1-C, UAS-trpA1-D*.^58^ D. Smith provided the *lush-GAL4*.^52^

### Chemicals

The following were from Sigma-Aldrich: benzaldehyde (B1334), propionic acid (81910), isovaleric acid (129542), citronellal (27470), AITC (W203408), and poly-L-ornithine (P4957). The sources of the following chemicals are as indicated: lemongrass oil (Nature’s Alchemy Pure Essential Oils), R-limonene (Fluka, 62122), DMSO (JT Baker, 67-68-5), and H2-DCFDA (Thermo Fisher Scientific, D399).

### DART2 assays

We constructed each assay tube by assembling one 150 mm-long Pyrex tube capped at either end with an odorant carrying cap. The odorant carrying caps were designed using Microsoft 3D Builder and printed using a Formlabs II 3D printer and black resin (FLGPBK01). The ‘Odorant-cap.stl’ 3D printer file is included in the supplementary information. The caps were washed in isopropyl alcohol for 1 hr., cured at 60 °C under violet light illumination in a Formlabs curing oven (FormCure), washed with an odorless detergent (Alconox, Cat. no. 1104-1), followed by a wash in ethanol, and then baked at 65 °C for 12-16 hours to remove residual odors. To ensure that there was no contact-chemosensation in the assay, the cap had meshed chambers to create an 8 mm-wide separation betweem the odorant and the flies in the assay tube.

We reared all flies at 25 °C. To prepare the flies for the assays, we collected flies 1-3 days post-eclosion from culture vials using CO_2_, placed flies in individual vials containing fresh food and allowed them to recover from the CO_2_ exposure for 48-72 hours. In preparation for the assay, the flies were transferred to each tube by attaching a 3D-printed custom funnel on the pyrex tube and tapping the flies into the tube. The ‘Funnel.stl’ 3D printer file is included in the supplementary information. We used Clear resin (Formlabs FLGPCL04) to print the funnel as we found that builds made out of clear resin offered better impact-resistance. The flies were then distributed homogeneously in each tube by gently tilting and tapping the tube. The flies were allowed to acclimatize in the tubes for 20 minutes. 5 μL of the solvent control and the test odor were added to filter paper rounds (Whatman cat no. 1001-6508), and then placed in the external chambers of the odor caps. The external chambers were then capped with lids broken off from 500 μL Eppendorf tubes. The tubes were immediately placed in a test chamber, similar to a previously described four field chamber,^76^ which we repurposed for these assays. The test chamber was maintained at 23 °C, and the temperature was monitored using a Thorlabs TSP-01 thermal probe. Flies were illuminated by an array of near infrared LEDs. Fly locomotion was captured using a CCD camera equipped with a 850 nm cutoff filter (Edmund Optics), at the rate of 1 frame/sec for 60 min. Fly movements were analyzed per-frame using a custom Python script. The transverse components of the average position of the flies or the center of mass (CoM) were calculated using CoM_tracker (code attached in supplementary text file “CoM_tracker”). The CoM per frame were calculated based on the formula: CoM = Σm_n_*x_n_/(M*L). We defined the terms as follows: m, the pixel count that crossed a preset brightness threshold to distinguish from the background in each video; x, the lengthwise coordinate of each count; M, the total number of counts; L, the length of the tube in pixels.

The avoidance index was calculated using PI_tracker (code attached in supplementary text file “PI_tracker”). We bisected each tube. Flies in the half of the tube that was nearest the solvent were considered to be repelled by the odor. Flies in the half of the tube nearest the side with the test odor dissolved in the solvent were considered to be attracted to the test odor. The avoidance index per frame was as follows: A.I. = (Σflies in odorant side - Σflies in solvent side)/Σall flies in tube.

The average CoM was obtained for the range of CoM values per frame when the behavior response reached a maximum followed by an approximately steady state. The response to each odor was unique and therefore the time windows to pick the average CoM were unique for every odor (Figure S7) ー propionic acid: 5 - 25 mins, R-limonene: 0 - 60 mins, AITC: 20 - 60 mins, citronellal: 20 - 60 mins, isovaleric acid: 5 - 25 mins, lemongrass: 20 - 60 min. All odorants were dissolved in DMSO, with the exception of propionic acid, which was dissolved in distilled H_2_O.

The odor caps and tubes were cleaned for reuse after every assay. The caps and tubes were first soaked in distilled H_2_O saturated with Alconox odorless soap for 1 hour. The soap was then completely washed off, the caps and tubes were soaked in 50% ethanol for 1 hr, and washed in distilled H_2_O. The caps and tubes were then baked for 20-24 hours at 65 °C.

Light intensities were measured with a Thorlabs PM100D power meter. LEDs were mounted on heatsinks and placed above the assay tubes such that there was uniform illumination of the entire tube length: UV (365 nm, 7021.365, Waveform lighting), blue (450 nm; Samsung LM561C), green (530 nm, 7041.525, Waveform lighting) and red (630 nm, 7041.630, Waveform lighting). The temperature inside the test chamber was monitored using a Thorlabs TSP-01 probe and maintained by attaching a 24 V exhaust fan at the top of the enclosure. The test chamber was vented at the bottom so that room air was able to flow in while stray light from the exterior was not permitted in.

### Immunohistochemistry

For reporter labeling of tissue, we used *UAS-GFP*, *UAS-RFP* and *LexAop-GFP* expressed under control of the indicated *GAL4* and *LexA* drivers at 25 °C. Flies were fixed in 4% paraformaldehyde (PF) in phosphate-buffered saline (PBS) for 30 minutes at room temperature (RT, ∼22 °C). The fixed animals were washed 6x with PBS containing 0.3% Triton X-100 (PBS-T) at RT. Antennae and brains were dissected in PBS-T. All tissue samples were washed 3x in PBS-T and blocked for 1 hr in PBS-T with 5% goat serum. The samples were incubated with primary antibodies diluted in blocking solution for 48 hours at 4 °C: mouse nc82 (1:100, Developmental Study Hybridoma Bank), chicken anti-GFP (1:1000, A-11122, Invitrogen), and rabbit anti-dsRed (1:1000, 632496, Clontech) or mouse anti-nc82 (DSHB). The samples were washed 3x for 20 min each in PBS-T and then incubated with secondary antibodies for 24 hrs at 4 °C: Alexa Fluor 488 goat anti-chicken(1:200, A-11039, Invitrogen), and Alexa Fluor 568 goat anti-rabbit (1:200, A-11011, Invitrogen) or Alexa Fluor 568 goat anti-mouse (1:200, A-11004, Invitrogen). The samples were washed 3x in PBS-T. Brain samples were mounted with Vectashield mounting media (Vectorlabs). Antenna and maxillary palp samples were mounted in Rapiclear (Sunjin Lab). All samples were imaged on an LSM 700 or LSM 900 confocal microscope (Zeiss). Images were prepared using Zen2 (Zeiss) or Fiji (ImageJ).

### Electroantennography

Flies were trapped in 200 μL tips and mounted under an inverted microscope (Nikon Eclipse FN1). A steady stream of humidified air was applied over the flies. The ground and recording electrode pipets were pulled using a Sutter P97 puller using 1B150F-3 borosilicate glass (Warner Instruments) to obtain a tip diameter that showed a resistance of 0.5 - 1 MOhms. Both the recording and reference pipettes were filled with electrolyte solution (108 mM NaCl, 4 mM NaHCO_3_, 1 mM NaH_2_PO_4_, 5 mM KCl, 2 mM CaCl_2_, 8.2 mM MgCl_2_, 5 mM HEPES, 10 mM sucrose, 5 mM trehalose). The pipet tips were coated with conducting cream (Sigma Creme, AD Instruments). The reference electrode was placed at the base of the antenna on the frons between the eyes, and the recording electrode was placed at the tip of the antenna. 5 μL of each odorant was applied to a filter paper round placed inside a 1 mL tip. Odor air puffs were applied for 500 ms through the 1 mL pipet tip positioned 1.5 cm from the fly, using a Syntech air pump. The responses were recorded using EAGPro (Syntech Oeckenfels GmbH) at the rate of 5kHz.

### DCFDA imaging

Antennae were dissected and placed in artificial hemolymph solution (AHLS) inside a 3D printer imaging chamber printed using black resin (Formlabs, FLGPBK01) (ImagingChamber.stl 3D printer file; see Supplementary Information). The imaging chamber had a thin perforation at the bottom that allowed a single antenna to be placed securely. Each antenna was then incubated for 45 min at RT with 2′,7′- dichlorofluorescein diacetate (DCFDA) at a final concentration of 2 μM in AHLS, washed 3x with ALHS, and imaged on a Zeiss LSM 900 confocal microscope. Baseline fluorescence at 530 nm was recorded for 30 sec before application of the UV stimulus (360 nm) by switching to the inbuilt DAPI filter in the LSM 900 for 1 sec. Following UV stimulation, DCFDA fluorescence was recorded every 3 sec for 2 minutes, and the steady-state peak fluorescence was used to compute the change in fluorescence from the baseline reading.

### Patch-clamp electrophysiology

S2 cells were cultured at RT in Schneider’s *Drosophila* medium (21720, Gibco) and transfected using either Lipofectamine 2000 (Invitrogen) or X-tremeGENE HP DNA Transfection Reagent (Roche) with the control plasmid (pAc5.1) encoding *GFP* only, or the plasmid encoding *GFP* and either *trpA1-C* or *trpA1*-*D*. The transfected cells were plated on poly-L-ornithine-coated 12 mm coverslips and assayed by whole-cell clamp 48 hours post-transfection. Drugs were dissolved in extracellular saline (140 mM NaCl, 5 mM KCl, 1 mM MgCl_2_, 2 mM CaCl_2_, 10 mM HEPES, 10 mM glucose), which was buffered to pH 7.4 with NaOH and with an osmolarity adjusted to 300 mOsm with mannitol. Cells were patched using borosilicate pipets (BF150-86-10, Sutter Instrument) with a pipet resistance of 5-7 MOhms. The pipet solution contained 140 mM CsCl, 1 mM MgCl_2_, 0.05 mM EGTA, 10 mM HEPES, which was buffered to pH 7.2 with CsOH and with an osmolarity adjusted to 290 mOsm with mannitol. Both the extracellular and pipet solutions were sterile filtered with 0.22 μm sterile filters. Drugs were added to cells using a gravity-driven perfusion system. Isolated cells were voltage-clamped in the whole-cell mode using an Axon 200B amplifier and 1440A Digitizer (Molecular Devices). Currents were recorded at 10 kHz. Cells were held at −60 mV with voltage ramps applied every 5 sec, which linearly increased from −80 to +80 mV within 200 ms to extract current vs. voltage characteristics. To test for membrane resealing, a membrane test pulse of −10 mV was applied for 20 msec within each voltage-clamp cycle 100 msec before application of the +80 to −80 mV voltage ramp. Cells that resealed were eliminated from analysis.

## Acknowledgments

This work was supported by grants to C.M. from the National Institute on Deafness and Other Communication Disorders (DC007864 and DC016278) and the National Institute of Allergy and Infectious Diseases (AI165575). We would like to thank Matthieu Louis, Angela Bontempo and David Tadres for technical suggestions.

## Declaration of Interests

The authors declare no competing interests.

## Figure legends

**Figure S1.**
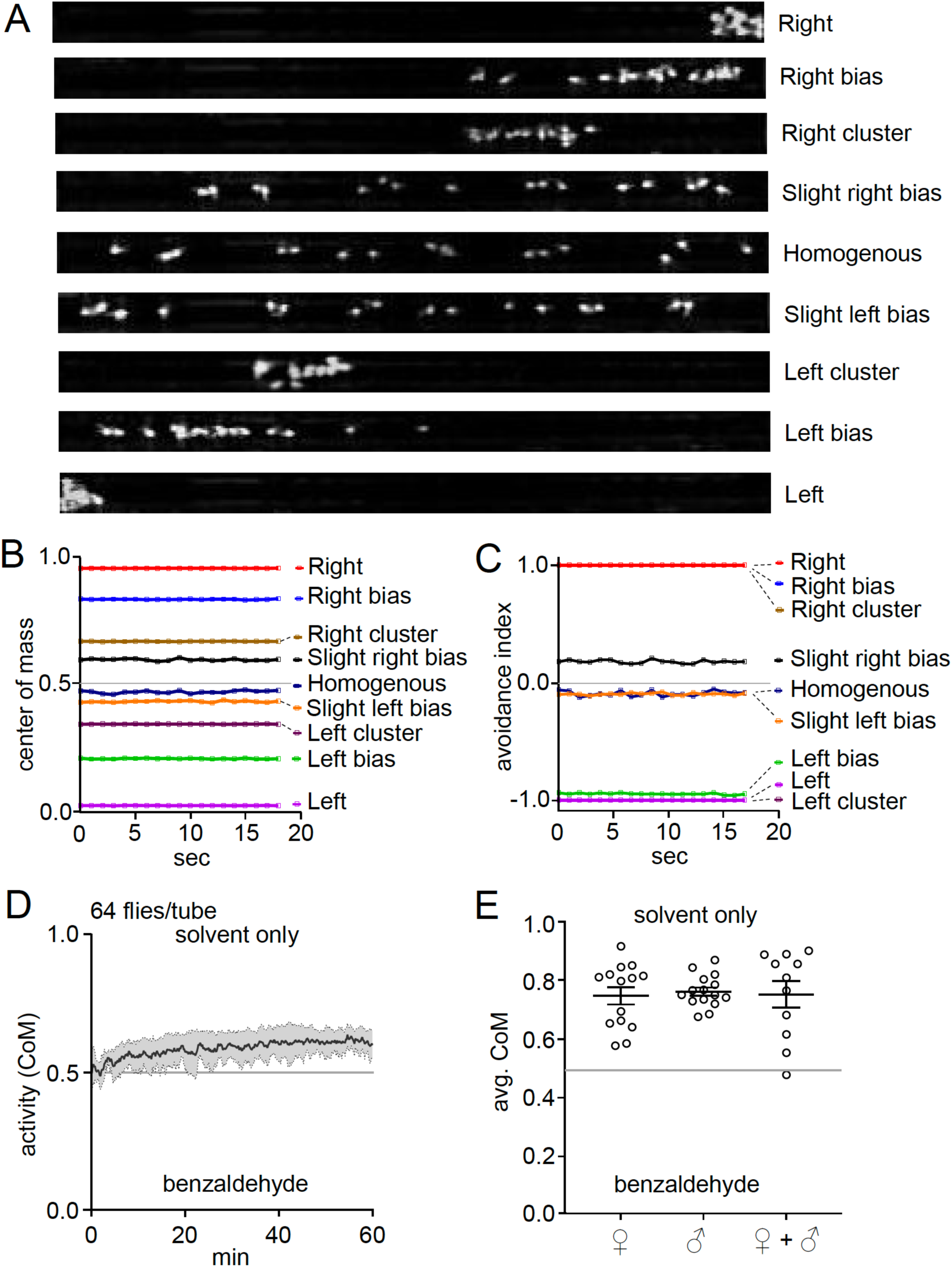
Design and calibration of the experimental paradigm for DART2 odor assay, Related to Figure 1. (A) Images of experimental tubes carrying flies immobilized by CO_2_ and arranged in 9 different positions simulating different outcomes observed in DART2 assays. (B) Tubes shown in (A) were analyzed using CoM as the measure of fly positions. See the Methods section (DART2 assays) for the CoM formula. (C) Tubes shown in (A) were analyzed using the avoidance index as the measure of fly positions. See the Methods section (DART2 assays) for the avoidance index formula. (D) Response to 1% BA with 64 flies per tube. (E) Flies do not show any sexual dimorphism in the two-way assay. Average CoM of control flies in response to 1% BA. Assay tubes contained males, females, or both. n = 11 – 15. ANOVA followed by Dunn’s multiple comparisons test. Error bars indicate mean ± SEMs. Differences were not significant.

**Figure S2.**
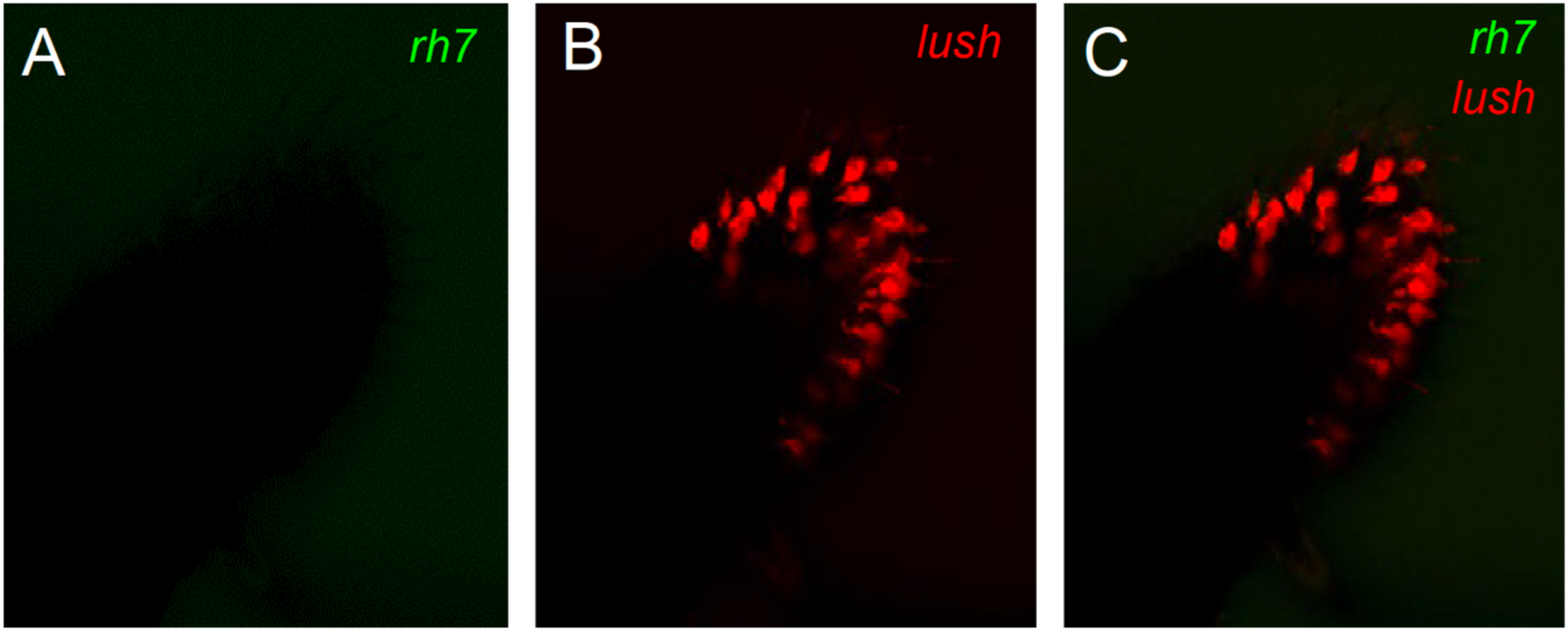
Testing for rh7 reporter expression in the antenna, Related to Figure 3. (A) Cross-section of antenna indicating lack of *lexAop-mCD8:GFP* expression driven by the *rh7-LexA*. The antenna was stained with anti-GFP. (B) Same antenna as (A) showing *UAS-mCD8::RFP* driven by the *lush-GAL4* and stained with anti-RFP. (C) Overlay of (A) and (B).

**Figure S3.**
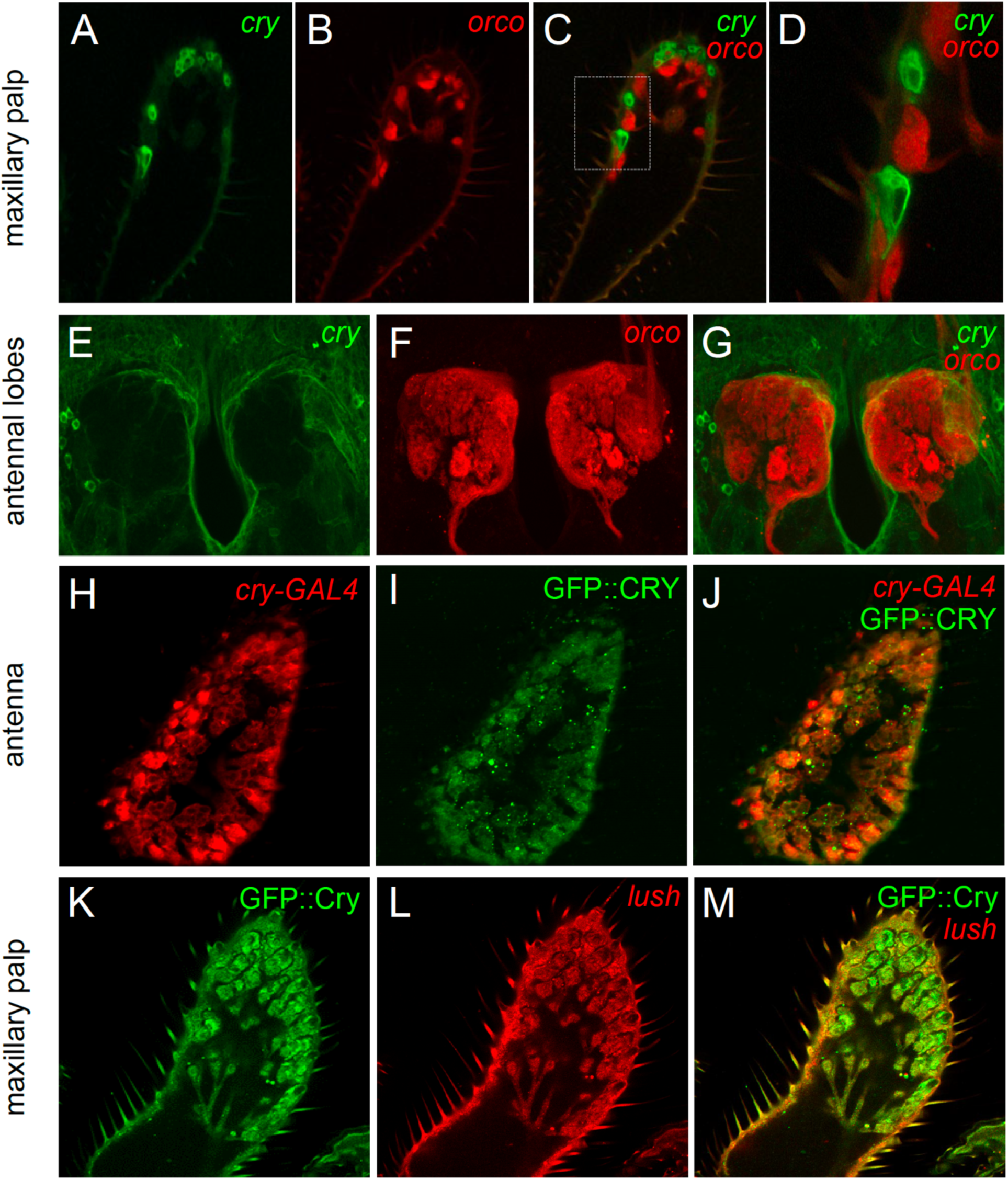
Expression of *cry*, *orco* and *lush* reporters in olfactory organs, Related to Figure 4. (A) Cross-section of maxillary palp showing *UAS-mCD8::GFP* expression driven by the *cry-GAL4* and stained with anti-GFP. (B) Same sample as (A) showing expression of *RFP* driven directly by the *orco* promoter (*orco-RFP*) and stained with anti-RFP. (C) Merge of (A) and (B). (D) Magnified section marked in (C) by white dotted rectangle. (E) Z-stack of antennal lobe showing *UAS-mCD8::GFP* expression driven by *cry-GAL4* and stained with anti-GFP. (F) Same sample as (E) showing *RFP* driven directly by the *orco* promoter (*orco-RFP*) and stained with an anti-RFP. (G) Merge of (E) and (F). (H) Antennal cross-section showing expression of *UAS-mCD8::DsRed* driven by the *cry-GAL4* and stained with anti-DsRed. (I) Same sample as (H) showing expression of GFP-tagged Cry (GFP::Cry) stained with anti-GFP. (J) Merge of (H) and (I). (K) Maxillary palp cross-section showing expression of GFP::Cry stained with anti-GFP. (L) Same sample as (K) showing expression of *UAS-mCD8::DsRed* driven by the *lush-GAL4* and stained with anti-DsRed. (M) Merge of (K) and (L).

**Figure S4.**
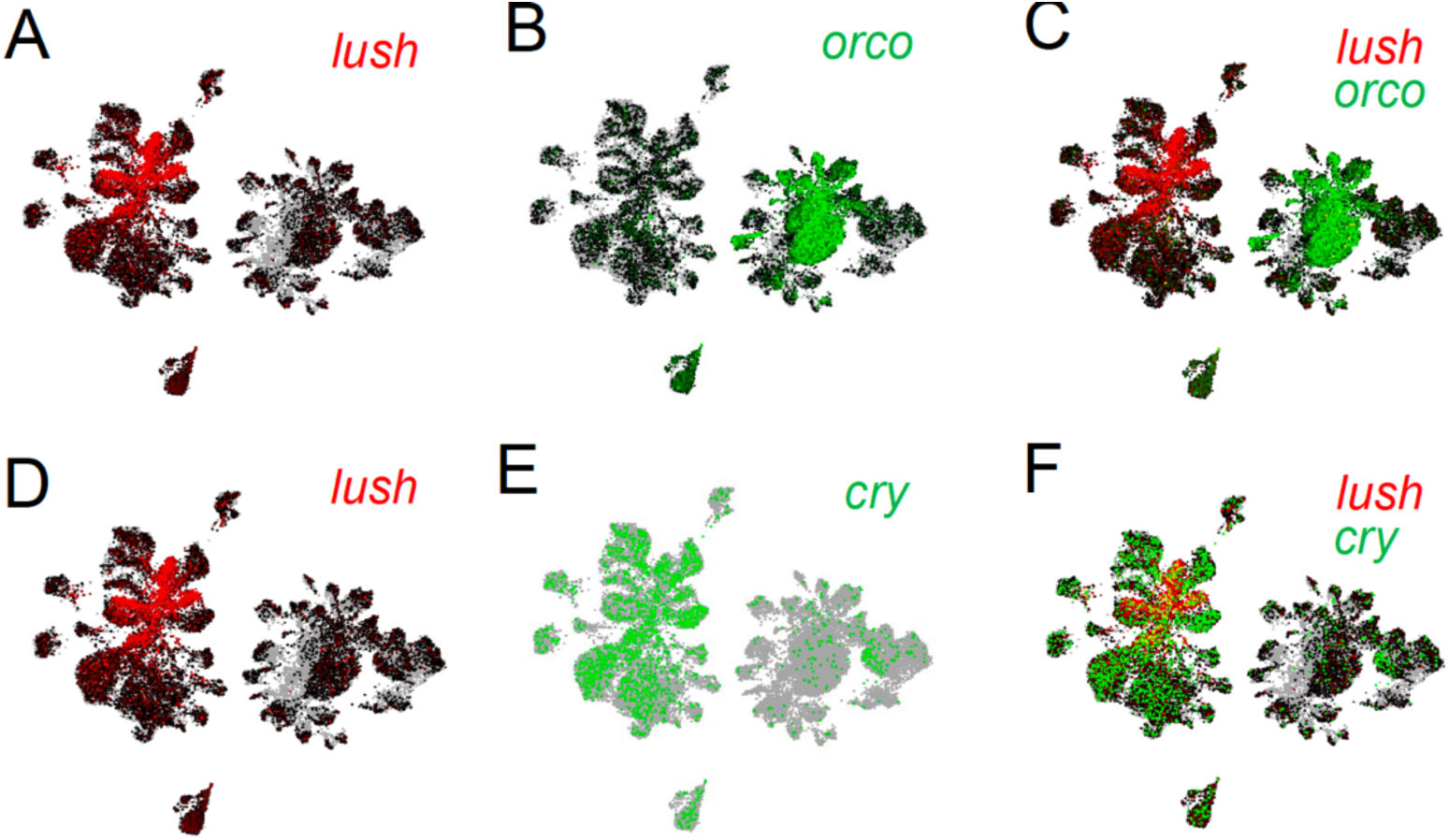
Abundance of *cry* transcripts in the antennae, Related to Figure 4. mRNA transcript levels plotted using FlyCellAtlas (https://flycellatlas.org/) and visualized using the Scope platform with the following method: SmartSeq2> Antenna> Stringent> s_fca_biohub_antenna_10X_ss2. (A) *lush* (red). (B) *orco* (green). (C) Merge of (A) and (B). (D) *lush* (red). (E) *cry* (green). (F) Merge of (D) and (E).

**Figure S5.**
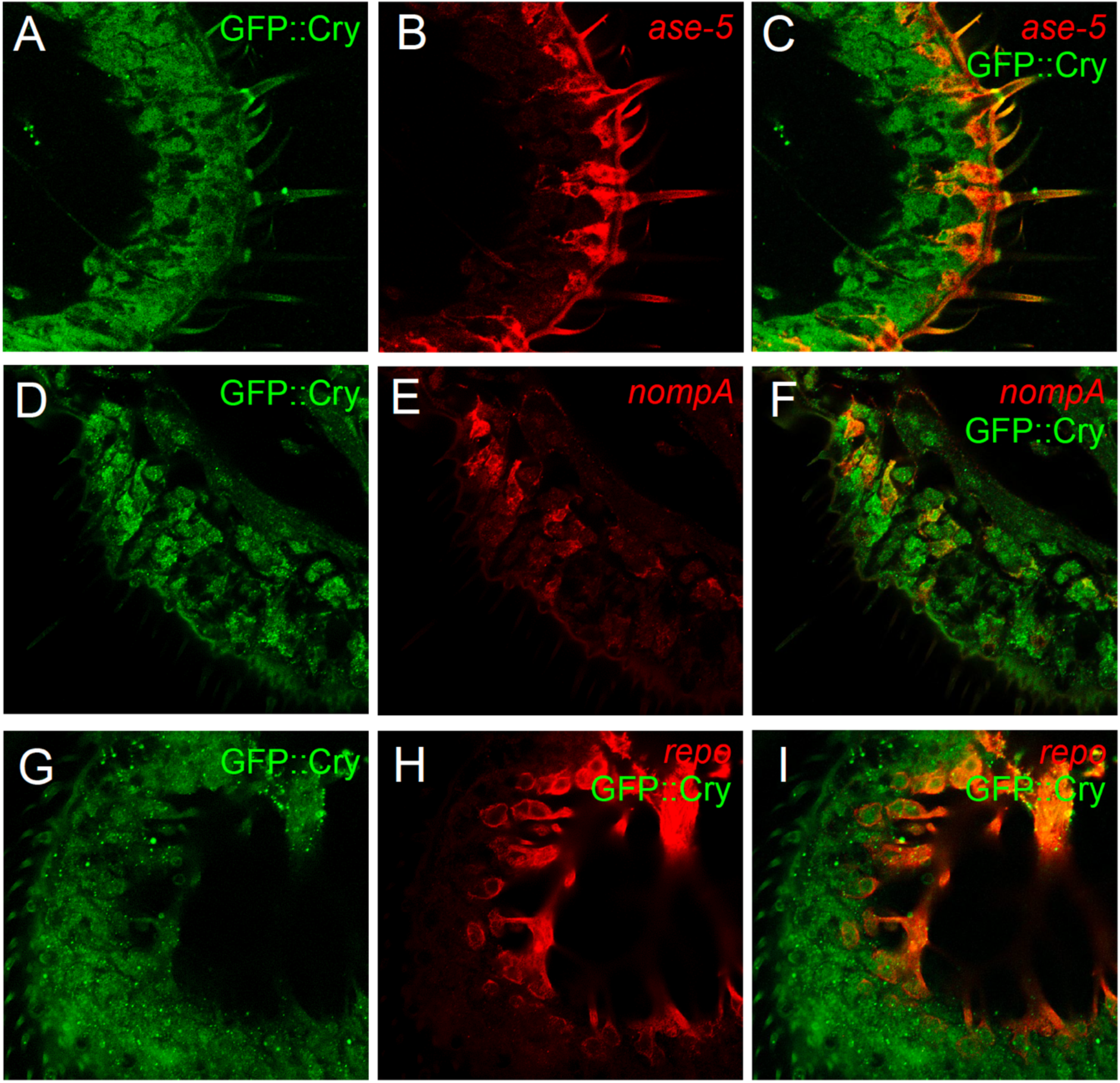
Testing for expression overlap in the antenna between the GFP::Cry reporter and support cell-type reporters, Related to Figure 4. (A) Cross-section showing expression of GFP::Cry stained with anti-GFP. (B) Same sample as (A) showing expression of *UAS-mCD8::DsRed* driven by the *ase5-GAL4* and stained with anti-DsRed. (C) Merge of (A) and (B). (D) Cross-section showing expression of GFP::Cry stained with anti-GFP. (E) Same sample as (D) showing expression of *UAS-mCD8::DsRed* driven by the *nompA-GAL4* and stained with anti-DsRed. (F) Merge of (D) and (E). (G) Cross-section showing expression of GFP::Cry stained with anti-GFP. (H) Same sample as (G) showing expression of *UAS-mCD8::DsRed* driven by the *repo-GAL4* and stained with anti-DsRed. (I) Overlay of (G) and (H).

**Figure S6.**
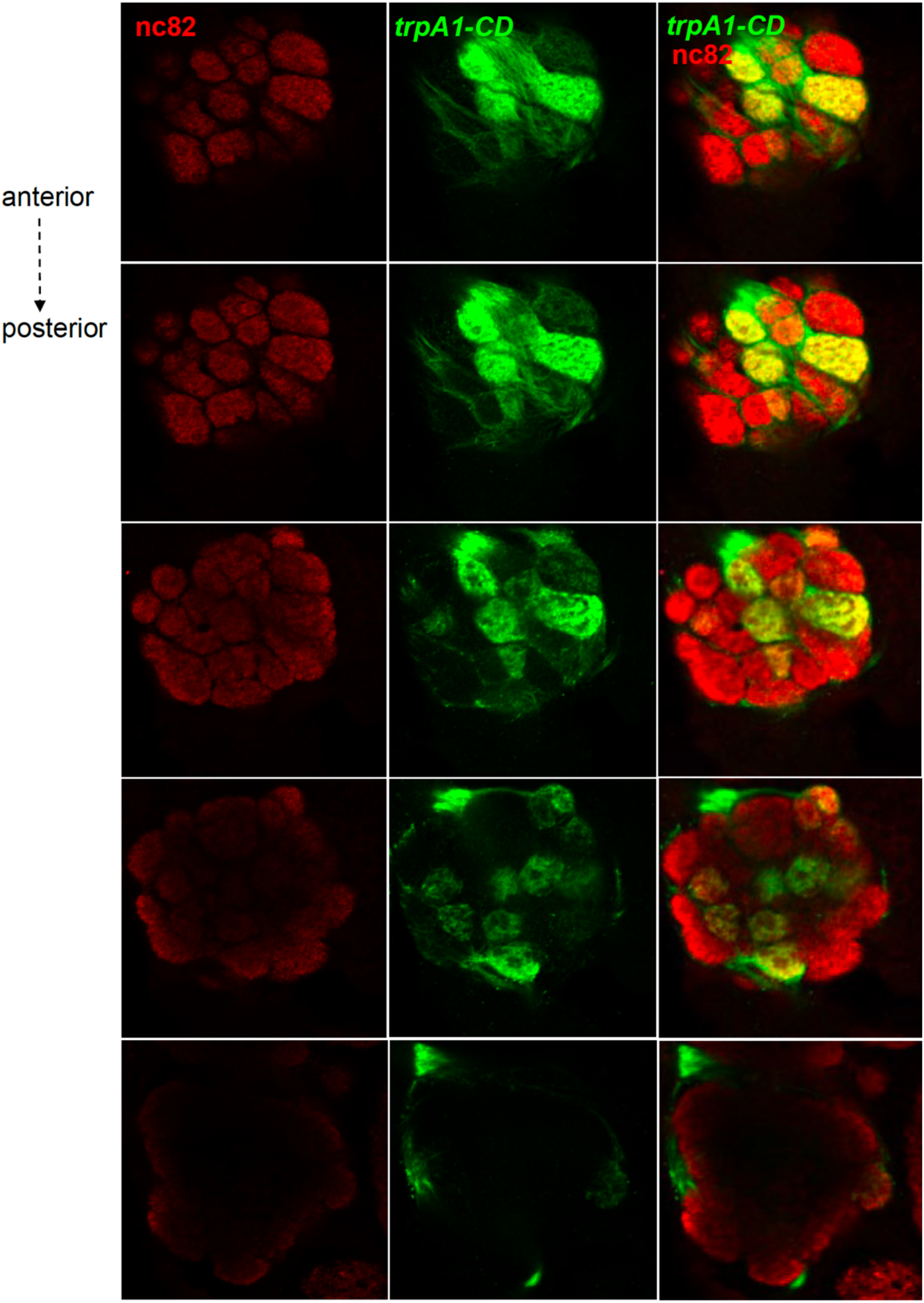
Cross sections of the antennal lobe showing *trpA1-CD* ORN processes, Related to Figure 6. *UAS-mCD8::GFP* was driven by the *trpA1-CD-GAL4* and stained with anti-GFP (green). The tissue was also stained with anti-nc82. The distance between every section and the previous one, starting from the second section was 1, 3, 5 and 7 μm respectively.

**Figure S7.**
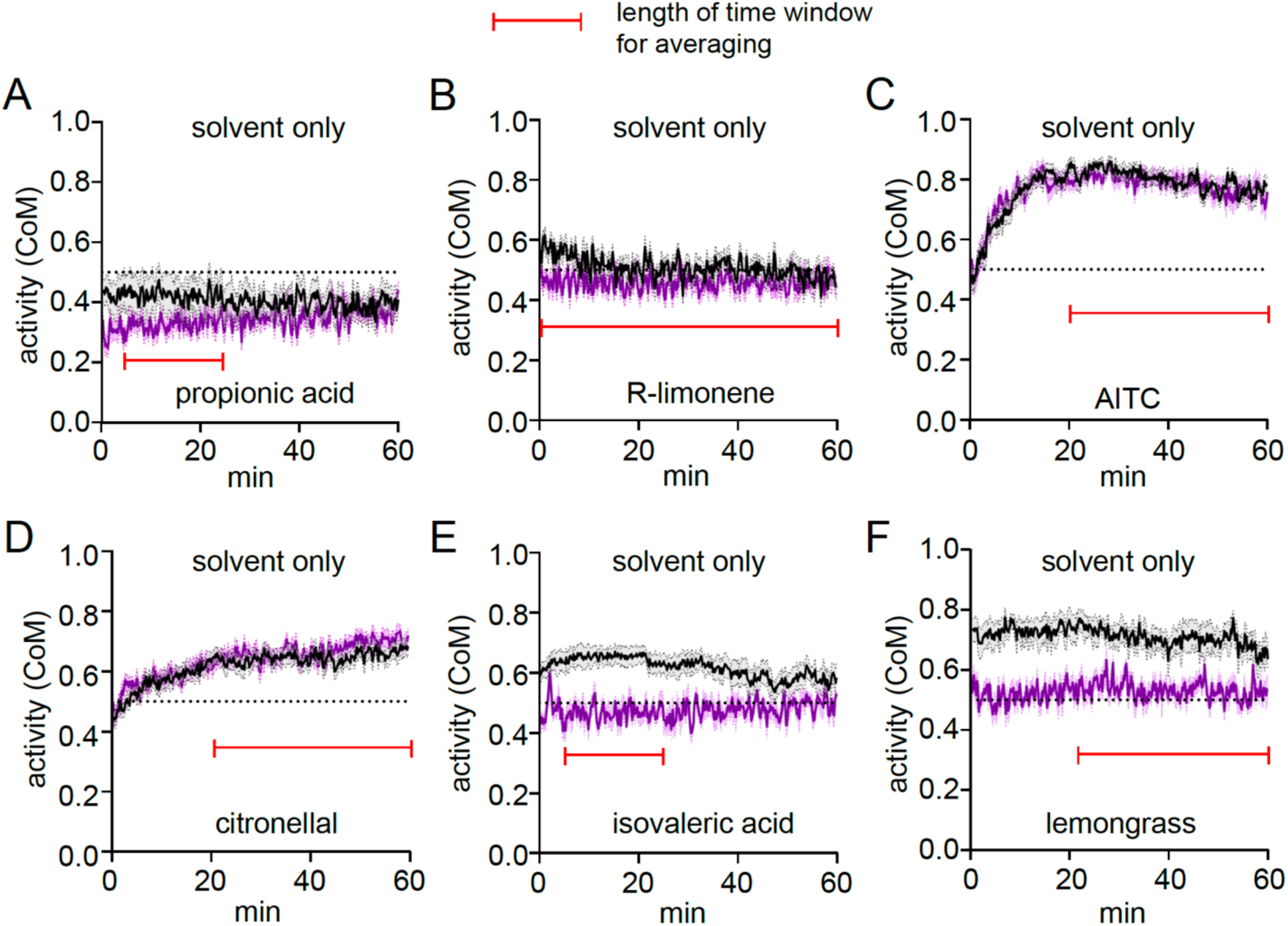
Time courses for behavior responses to odorants in DART2 assay, Related to Methods. The various odorants used are indicated below. The time-window that was used to calculate the average CoM is indicated by the red bracketed lines. (A) 1% propionic acid, (B) 1% R-limonene, (C) 0.1% AITC, (D) 3% citronellal, (E) 0.01% isovaleric acid, (F) 3% lemongrass.

## Notes

### Competing Interest Statement

The authors have declared no competing interest.

